# A multi-modal flow phantom for quantitative PET/Spectral CT

**DOI:** 10.64898/2026.06.15.731705

**Authors:** Elizabeth J. Li, Sophie Lammers, Yinglin Ge, Stephen McDonald, Michael Geagan, Joshua Scheuermann, Austin R. Pantel, Peter B. Noël, Joel S. Karp

## Abstract

**Purpose:** In this work, we aimed to establish a flow phantom for multi-modal PET and spectral CT imaging to improve blood flow quantification.

**Methods:** A modular flow phantom was built with materials compatible with both PET and spectral CT. A peristaltic pump was used to allow for recirculation. Pores were installed through the aorta to allow for tissue exchange between the blood and tissue compartments, and valves were placed in line with the aorta to control the pressure gradient between compartments. We characterized the system using saline bolus experiments, dynamic PET imaging, and iodine-based spectral CT acquisitions. A blood flow (K1) of 1.0 mL/min/mL with a pressure range of approximately 1.0-3.0 psi was targeted. Using compartmental modeling, we estimate K1 across phantom configurations and evaluate the consistency of perfusion-related parameters derived from saline, PET, and spectral CT measurements.

**Results:** With the four pore, two valve configuration, target K1 of 1.0 mL/min/mL was achieved with a physiologic pressure range (2.2-3.5 psi) and a pump speed of 150 rpm. Further, the flow phantom was also able to recapitulate K1 across a range of values through adjustable modifications to the phantom configuration.

**Conclusions:** We present a modular multimodal flow phantom with a tissue-mimicking compartment, vascular tubing with an adjustable number of pores and valves, and 3D-printed components to support tunable exchange between blood-pool and tissue compartments and controlled dynamic perfusion imaging with same-session PET and spectral CT. Such a setup will enable the development of multi-modal approaches for evaluating tissue perfusion.

## Introduction

Quantitative assessment of tissue perfusion is central to many biomedical imaging applications, including oncologic characterization [1–4], cardiovascular disease assessment [5, 6], and therapy monitoring [2, 7, 8]. Dynamic imaging enables estimation of tracer delivery and tissue exchange, providing physiologically meaningful parameters such as blood flow and transport rates. In positron emission tomography (PET), compartmental modeling approaches are well established for quantifying tracer kinetics and estimating parameters that describe transport from the vascular compartment into tissue [9, 10]. In parallel, advances in spectral computed tomography (CT) enable material-specific imaging and quantitative mapping of contrast agents such as iodine [11], creating opportunities to estimate perfusion-related parameters from CT-based dynamic acquisitions.

PET and spectral CT provide complementary information for dynamic perfusion imaging. PET offers high sensitivity to radiotracer quantification and mature kinetic modeling frameworks [9, 10, 12], whereas spectral CT provides high spatial resolution and quantitative iodine concentration maps [11, 13, 14]. As long-axial-field-of-view PET systems and quantitative spectral CT techniques become increasingly available, including the first, and to our knowledge, only integrated PET/spectral CT platform developed at the University of Pennsylvania, there is growing interest in coordinated PET/CT approaches for evaluating tracer delivery, tissue perfusion, and contrast-agent kinetics. However, direct comparison of PET-derived and CT-derived perfusion metrics remains challenging because the modalities differ in acquisition physics, temporal sampling, spatial resolution, tracer delivery, and contrast-agent behavior.

A major limitation in developing and validating quantitative perfusion imaging methods is the lack of controlled experimental reference conditions. In vivo measurements are affected by physiologic variability, tracer-specific extraction, motion, blood input-function uncertainty, and the absence of an independent reference standard for local tissue flow. Conventional imaging phantoms [15, 16] are widely used for scanner calibration and image-quality assessment [17], but most are static or optimized for single-modality performance testing. Dynamic flow phantoms have been developed for selected PET, SPECT, CT, and cardiac applications [18–22], yet there remains a need for a practical, reproducible platform that supports adjustable flow rates, multimodal imaging compatibility, and kinetic modeling of both radionuclide and iodine bolus measurements.

Such a platform must meet several technical requirements. It should generate stable and controllable flow conditions over a physiologically relevant range, demonstrate reproducible temporal bolus dynamics for kinetic analyses, and permit systematic variation of experimental parameters such as pump speed, pressure, and exchange between vascular and tissue-mimicking compartments. Adequate mixing within the tissue compartment is required for interpretable time-activity or time-concentration curves, while the geometry and composition of materials must remain compatible with both PET and spectral CT. These requirements are particularly important for evaluating whether PET- and iodine-based spectral CT measurements produce consistent perfusion-related parameters under matched experimental conditions.

In this work, we present a modular multimodal flow phantom for controlled dynamic perfusion imaging with PET and spectral CT. The phantom uses a tissue-mimicking compartment, vascular tubing with an adjustable number of pores and valves, and 3D-printed components to support tunable exchange between blood-pool and tissue compartments. We characterize the system using saline bolus experiments, dynamic PET imaging, and iodine-based spectral CT acquisitions. Using compartmental modeling, we estimate K1 across phantom configurations and evaluate the consistency of perfusion-related parameters derived from saline, PET, and spectral CT measurements.

## Methods

### Phantom design

The phantom set-up is shown in Figures 1A and 1B, where parts shown in red were 3D-printed. A 10.2 cm inner diameter cylinder (referred to as tissue space) was placed inside a NEMA torso phantom shell (Data Spectrum Corporation, Durham, North Carolina). To mimic the cross-sectional dimensions of the aorta [23], a 2.54 cm inner diameter soft Tygon tube was cut into two segments and attached to the ends of a 3D-printed curved pipe along the center line of the cylinder, which forms a U shape when assembled. This U is referred to as the blood pool or aorta.

**Figure 1.**
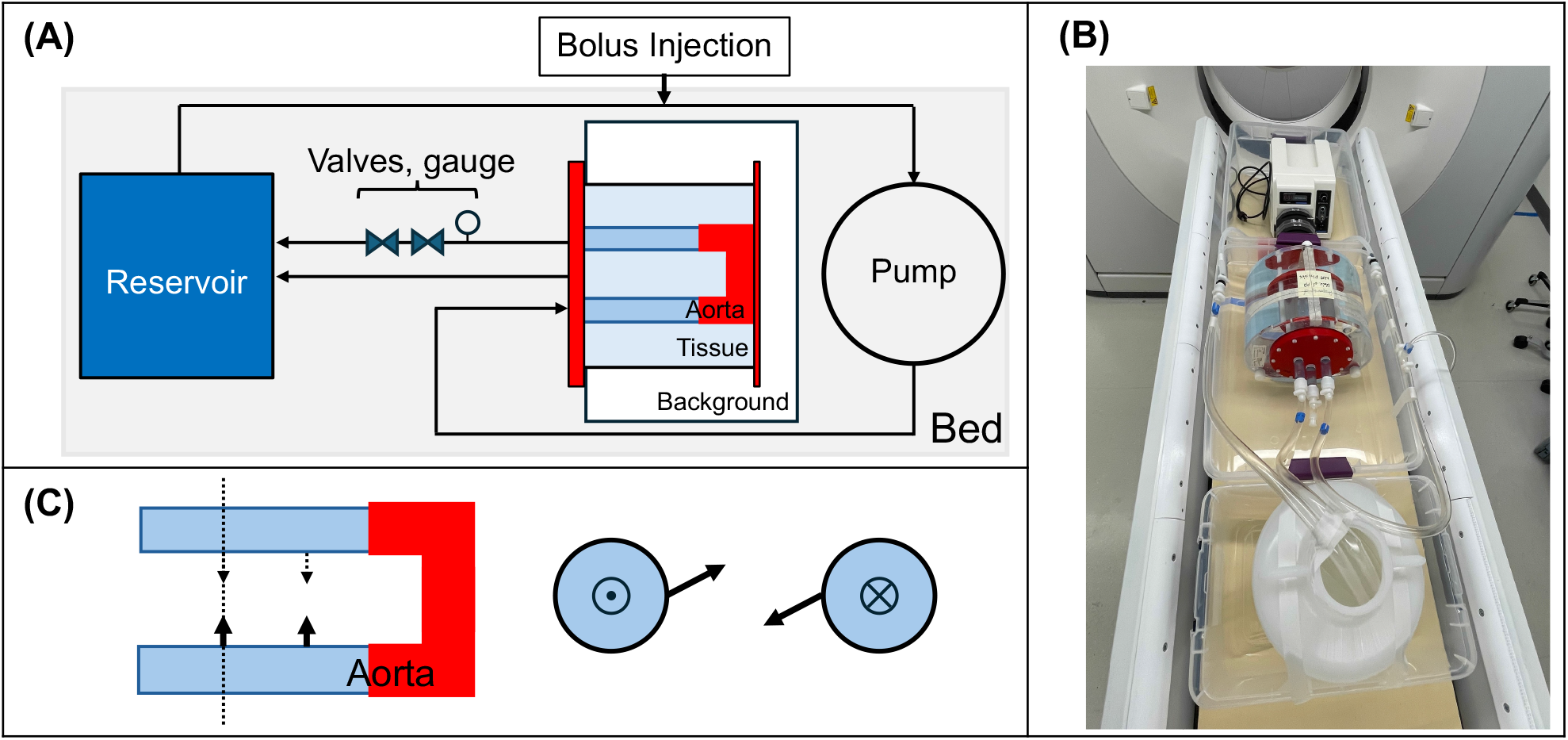
(**A**) Schematic of PET/CT imaging setup. 3D-printed parts are shown in red. When activated, the pump circulates water into the phantom, through the pores in the aorta (Tygon tubing), and back into the reservoir. (**B**) Experimental set-up on the PET/CT bed. The background compartment has been filled with water and the reservoir filled with approximately 5 liters of water for recirculation. (**C**) Inset of aortic tubing shows placement of pores (in this example 4 pores) in the coronal (left) and axial cross section (right) at the dotted line. Pores were placed symmetrically across the U of the aorta but were offset to allow for improved mixing in the tissue compartment.

Fluid exchange from the aorta to the tissue was accomplished across a wide range of flow rates through the introduction of pores to the walls of the aortic tubing. Zero, two, three, or four, 3-mm diameter pores were bored through identical but separate aortic tubing segments so that the number of pores, and thus the flow rate, could be changed as needed. Due to the flexibility of the tubing, attempts at using a smaller diameter drill bit resulted in pore collapse. Pore pairs were placed symmetrically across, and were evenly spaced along the two installed segments. When three pores were installed, a single central pore was installed on one segment, while two were installed in the opposite segment. For the even numbers of pores, pore pairs were angled anti-parallel from each other when installed (Figure 1C) to allow for increased mixing in the tissue compartment. Additionally, to further control flow rate from the aorta to the tissue compartment, one or two valves were added in-line with the aorta. The valves increased the pressure gradient between the aortic and tissue spaces, increasing flow to the tissue compartment.

Water-tight ports were installed in the 3D-printed lid and connected to tubing leading to and from the recirculation reservoir (Figure 1A). This allowed the solution to be recirculated within the blood pool via a peristaltic pump (WT600-2J, KZ25 pump head Langer Instruments, Tucson, Arizona) while maintaining a water-tight seal between the cylinder, torso shell background, and exterior of the torso phantom. 1.3 cm inner diameter tubing was used to circulate fluid throughout the system. The outlet from the tissue compartment was also inserted into the recirculation reservoir. An injection port was inserted in-line between the pump and the phantom.

### Benchtop calibration studies – volumetric flow

Benchtop studies were performed using a single-pass configuration, where tracer recirculation was eliminated by exclusion of the recirculation reservoir. For the initial calibration of volumetric flow in the aorta (units mL/min) versus pump speed, no pores were included in the aortic tubing, so that there was no tissue uptake from the blood pool. Pump speed was increased in increments of 50 rotations per minute (rpm, range 100-400 rpm), and the output volume from the aorta was recorded over time to calculate volumetric flow.

Volumetric flow was measured with additional modifications to the phantom setup, including changes in pump speed, pressure, and phantom configuration (e.g., number of pores, valves). With the four pore, two valve configuration, the relationship of pressure and volumetric flow was further assessed over a wide pressure range (0.3-2.65 pounds per square inch, psi) while pump speed was kept constant (150 rpm).

Additional benchtop studies included tissue exchange between the aorta and tissue space through the incorporation of pores (Figure 1C).

### Benchtop calibration studies – K1

After establishing the relationship of volumetric flow with pump speed and pressure, three sets of benchtop saline and food dye bolus studies were then performed, as a proxy for measuring flow without radioactivity during calibration studies. For saline studies, 10 mL of saline (approximately 90 parts per thousand, ppt) was administered as a bolus and the salinity of the tissue and aortic outputs were measured over time. Green food dye was added to the bolus to visually confirm mixing in the tissue compartment. Concentration flow, also called perfusion or K1 (mL/min/mL) was calculated as detailed in the kinetic modeling section.

Two of the three saline bolus studies involved evaluation of flow with variable pump speed and pressure with the four pore, two valve configuration. First, pump speed was altered in 50 rpm increments (range 100-300 rpm) and the valves were adjusted to maintain a physiologic range of 1.0 to 3.0 psi. For the second series of saline bolus studies, the valves were calibrated to obtain a target pressure of 2.3 psi (120 mm Hg) at 150 rpm. Pump speed was adjusted to 100, 150, and 200 rpm. Psi was recorded and K1 was estimated.

A third and final set of saline bolus studies was performed to estimate K1 across the possible pore and valve configurations. Pump speed was set to 150 rpm while the number of pores and valves were varied. The valves were adjusted to obtain a range of approximately 2.3-3.5 (120-180 mm Hg). K1 was estimated for each experimental setup.

### Phantom imaging studies

After benchtop calibration and saline bolus studies, PET and CT acquisitions were performed using the PennPET Explorer [24] and a CT with a 4-cm dual-layer spectral detector (IQon Spectral CT, Philips Healthcare, Eindhoven, Netherlands). After a non-contrast enhanced CT for attenuation correction, rubidium-82 (^82^Rb) injected via generator (CardioGen-82, Bracco Diagnostics, Milan, Italy) during a 7-minute PET acquisition. A serial, iodine-enhanced CT was then performed. The CT autoinjector (MEDRAD Flex CT, Bayer, Leverkusen, Germany) was loaded with a 1:10 iodine contrast (ISOVUE 370, Bracco Group, Milan, Italy, 370 mg iodine/mL):saline solution. PET and CT studies were performed in the same imaging sessions (Figure 1C). PET images were reconstructed with 4 × 4 × 4 mm^3^ voxels. Additional bolus and acquisition parameters are included in Table 1. A K1 of 1.0 mL/min/mL was targeted with a pump speed of 150 rpm. The valves were adjusted to maintain a physiologic pressure range of 1.2-4.0 psi (62 – 207 mm Hg). Most phantom imaging studies were performed with two configurations: two pores and one valve, and with four pores and two valves for finer control of the aortic pressure. These two configurations represent the extremes of the benchtop pore and valve configurations.

**Table 1:**
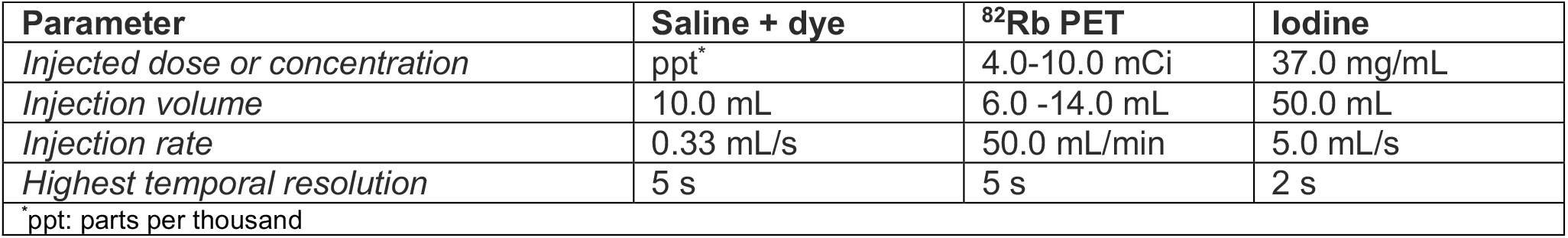
Phantom imaging study parameters.

| **Table 1: Phantom imaging study parameters** |  |  |  |
| --- | --- | --- | --- |
| **Parameter** | **Saline + dye** | **<sup>82</sup>Rb PET** | **Iodine** |
| *Injected dose or concentration* | ppt* | 4.0-10.0 mCi | 37.0 mg/mL |
| *Injection volume* | 10.0 mL | 6.0 -14.0 mL | 50.0 mL |
| *Injection rate* | 0.33 mL/s | 50.0 mL/min | 5.0 mL/s |
| *Highest temporal resolution* | 5 s | 5 s | 2 s |
| *ppt: parts per thousand |  |  |  |

### K1 estimation, kinetic modeling

For saline studies, flow phantom aortic and tissue outputs were manually collected over time. A salinity meter (Oakton Instruments) was used to record the time activity curves (TACs) for each region in units of ppt from the manually-collected samples.

Spectral CT iodine density maps (i.e. iodine, no water) were generated and quantified for all CT results using the vendor provided platform. PET and spectral CT image analyses were performed in MIM (MIM Software, Inc.). For the flow phantom, a 1-cm diameter, 3 cm in height cylindrical volume of interest (VOI) was placed in the aorta and tissue spaces on the ^82^Rb-PET and iodine CT images to generate image-derived input functions and tissue TACs for kinetic modeling. Due to under-mixing of iodine in the tissue compartment, two tissue VOIs were placed in areas of high and low iodine concentration and averaged for the kinetic modeling of iodine. The units for saline, ^82^Rb, and iodine TACs are mg/mL, kBq/cc, and mg/mL, respectively.

A 1-tissue compartment model [9] implemented in PMOD (Bruker Scientific LLC, Billerica, Massachusetts) was used to estimate K1 (units mL/min/mL), also referred to as concentration blood flow or tissue perfusion. Delay of bolus arrival was included in the fitting process. Blood volume fraction was fixed to zero.

### Statistics

To assess the overall behavior of the phantom with increasing pump speed, Pearson’s correlation was calculated between reference flow values from the manufacturer [25] and aortic volumetric flow (units mL/min). In this initial scenario, no tissue exchange was included in the set up (i.e., zero pores were introduced). After adding pores, the relationship between saline K1 and pressure with pump speed was also assessed via Pearson’s r. All statistical tests were performed in MATLAB (MathWorks Inc., Natick, Massachusetts)

## Results

### Phantom calibration

Figure 2 shows the results for the phantom calibration studies. There was a strong agreement between pump speed and aortic flow (Figure 2A) when no pores were present (Pearson’s r = 0.98, p = 0.002). This relationship was maintained when two pores were added (Pearson’s r = 0.95, p = 0.01). However, with the addition of valves, flow in the aorta decreased, with a corresponding increase in tissue flow, as expected. Figure 2B shows this relationship with four pores and two valves at 150 rpm, where the volumetric flow increased in the tissue as the valves were gradually closed and pressure in the aorta was increased. When pump speed was increased and a pressure of approximately 2.5 psi was maintained with manual adjustments to the valves, K1 plateaued (Figure 2C). When the valves were left unchanged, both K1 (slope: 0.4, r = 0.95) and psi (slope: 1.3, r = 1.0), increased linearly with pump speed. Together these results indicated that flow was less sensitive to pump speed and more sensitive to the pressure gradient between the aortic and tissue compartments. Further studies focused on a pump speed of 150 rpm.

**Figure 2.**
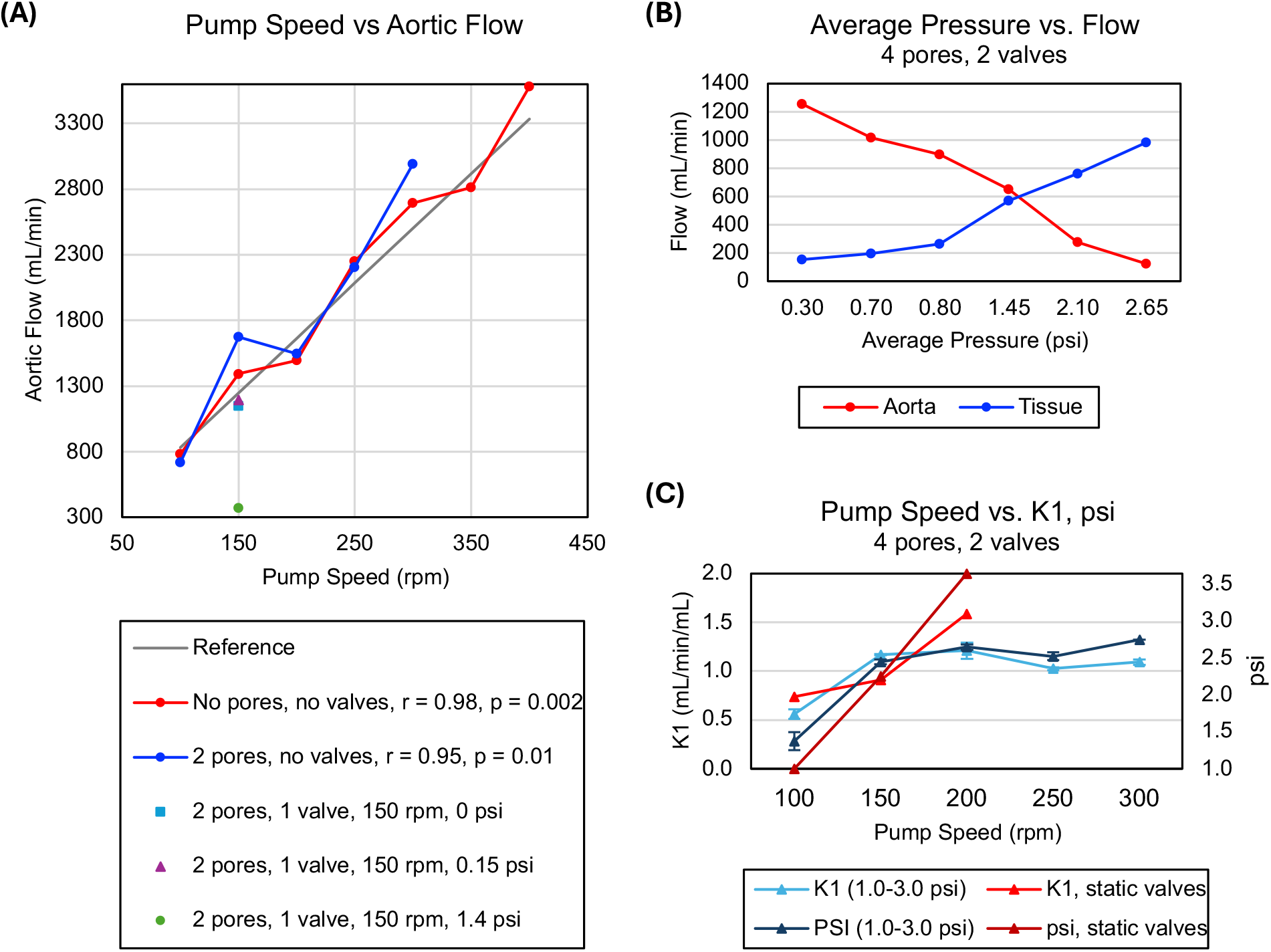
(**A**): Calibration of phantom setup, where flow in the aorta with various configurations was compared with known reference from manufacturer. (**B**): Flow versus pressure using 4 pores and 2 valves at 150 rpm. As expected, increased pressure on the aortic output resulted in higher flow to the tissue. (**C**): Increases in pump speed resulted in a plateauing of K1 and pressure with the 4 pore, 2 valve configuration when psi was maintained in a range of 1.0 – 3.0, but showed a wider range when valves were fixed in a static position.

### Flow estimates

Following the saline studies, ^82^Rb PET and spectral CT iodine mapping was performed for multiple configurations. In Figure 3, cross sectional images of the 2 pore, 1 valve configuration are shown. Since the PET and CT bolus injection rates, injected volumes, and temporal framings differ, the injected activity traverses the aortic space with different speeds, as evident from the faster washout and smaller FWHM of iodine (9.8 s) relative to ^82^Rb (8.0 s).

**Figure 3.**
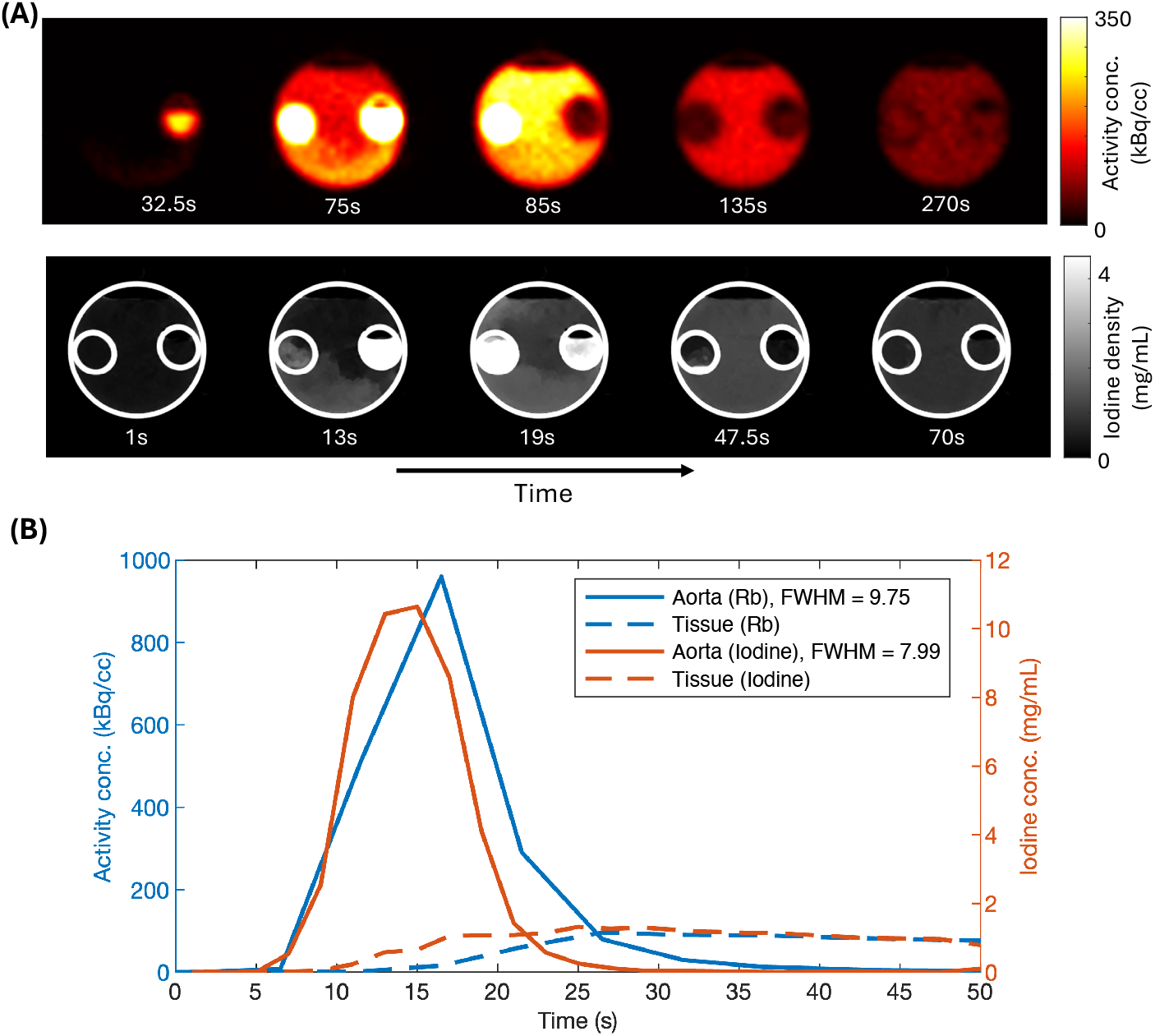
(**A**): Axial cross-sections of ^82^Rb PET (top) and iodine concentration images (bottom), showing the introduction of the tracers in the aorta, followed by the tissue space over time with the 2 pore, 1 valve configuration. (**B**): Representative early time activity curves for ^82^Rb (blue) and iodine (orange). The curves have been shifted and aligned in time to better compare aorta curves.

K1 was estimated for the various phantom configurations with saline (Figure 4), yielding a range of K1 from approximately 0.6 to 1.4 mL/min/mL across configurations with a pressure of 2.2 to 3.5 psi (114-180 mm Hg). Focusing on the two pore, one valve and four pore, two valve configurations, saline K1’s of 0.58 ± 0.07 and 1.04 ± 0.04 mL/min/mL, respectively were achieved. K1 values for ^82^Rb for the two pore, one valve and four pore, two valve configurations were 0.66 ± 0.02 mL/min/mL and 0.95 ± 0.14 mL/min/mL, respectively. For iodine studies, K1 was 0.72 mL/min/mL and was 0.86 ± 0.04 mL/min/mL for the two pore, one valve and four pore, two valve configurations, respectively. Overall, K1 was lower with ^82^Rb PET and iodine mapping as compared to the saline studies (Figure 4B). For all three bolus types, K1 was closer to the target of 1.0 mL/min/mL with the four pore, two valve configuration.

**Figure 4.**
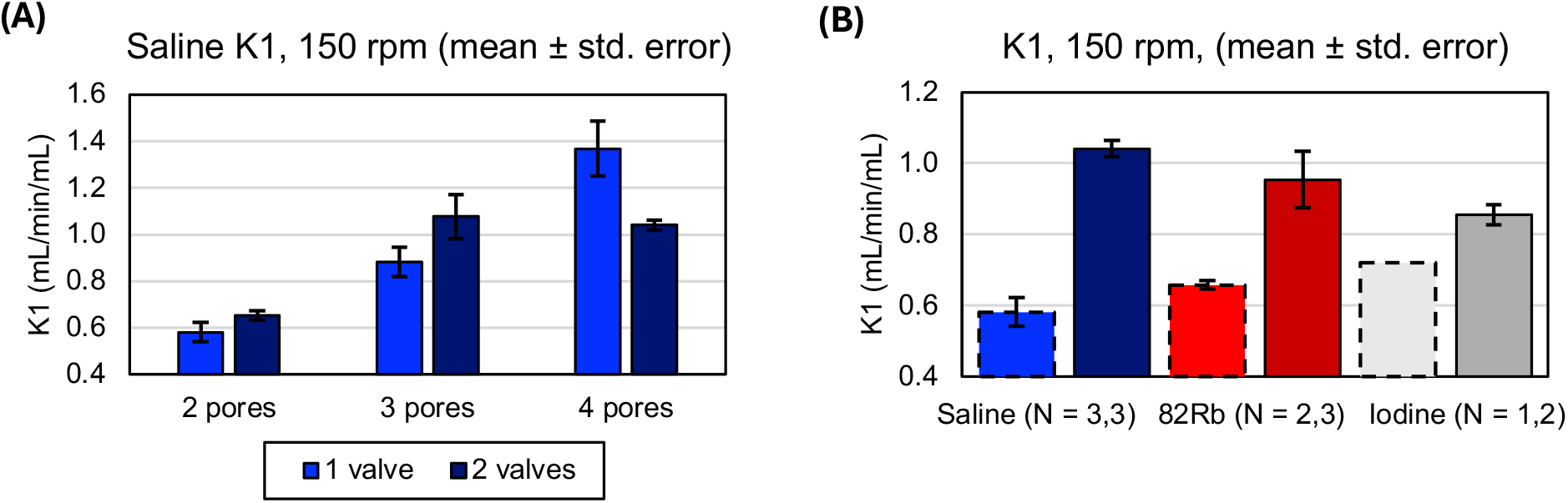
(**A**): 1-tissue compartment modeling estimates of K1 for various benchtop saline setups. Pump speed was held constant at 150 rpm and valves were adjusted to maintain a pressure range of 2.2 to 3.5 psi. (**B**): Based on results in (A) and Figure 2, 2 pore, 1 valve (dotted outlines) and 4 pore, 2 valve (solid outlines) configurations were used to estimate K1 with saline (blue) ^82^Rb (red) and iodine CT (gray). Fewer iodine studies were acquired, as evidenced by the horizonal labels.

## Discussion

We designed and built a dynamic, multimodal flow phantom for PET and spectral CT. The setup is capable of linear changes in volumetric flow with changes in pump speed. Finer control of flow was achieved by using valves to induce a pressure gradient across the aorta-tissue boundary. With the introduction of pores, the pressure gradient from the aorta to the tissue space impacted flow to a greater extent than pump speed. Across configurations, K1 was similar between iodine-enhanced dual-energy CT and PET-based measurements.

The spectral CT used in this work results in an axial field of view of 4 cm. To fully capture in vivo blood flow in organs such as the myocardium, these methods require either further technological development, either large z-coverage scanner configurations, or high-pitch helical acquisitions. Their development and validation will leverage this phantom platform, which supports calculation of K1 and enables, for the first time, multimodal imaging with multiple tracers without radiation dose deposition.

There are some limitations to the phantom design. The phantom does not allow for changes in first-pass extraction, which is both tracer- and flow-dependent *in vivo*, nor does it perfectly mimic human anatomy. However, our target parameter K1 represents a combination of extraction and flow, allowing for overall estimation of this parameter. Additionally, the phantom allows for controlled measurement of K1 with a flexible, benchtop-friendly and reproducible setup that is less subject to partial volume effects that might be seen in more anatomically realistic phantoms. Further, the Tygon tubing was selected with the diameter of the aorta in mind, such that this work is translatable to other PET scanners regardless of AFOV, but is particularly suited for LAFOV PET, where the descending aorta is always in the field of view of the scanner. With a wide range of possible K1 values (0.58 to 1.4 mL/min/mL) across configurations, this phantom can be used to mimic blood flow to a number of organs, such as gray matter, liver, and myocardium, which all have K1 values within this range, as measured with ^18^F-FDG and ^11^C-butanol [26, 27].

Future directions of this work will include narrowing of the aorta tubing to allow for variable blood vessel size, and refining pressure control and modification of the tissue compartment volume to achieve higher K1 values for tissues such as the kidney. There was also delayed mixing of the iodine contrast in the tissue compartment. The impact of circulating water temperature, which was not controlled here, as well as the strategic positioning and angle of the pores, will be investigated.

## Conclusions

We present a modular multimodal flow phantom for controlled dynamic perfusion imaging with same-session PET and spectral CT. While a target K1 of 1.0 mL/min/mL was achieved with a physiologic pressure range (2.2-3.5 psi) and a pump speed of 150 rpm, the flow phantom was also able to recapitulate K1 across a range of values through adjustable modifications to the phantom configuration. This phantom will enable the development of multi-modal approaches for evaluating tissue perfusion.

## References

1. Kim, S.H., A. Kamaya, and J.K. Willmann, CT Perfusion of the Liver: Principles and Applications in Oncology. Radiology, 2014. 272(2): p. 322–344.

2. van Dijken, B.R.J., et al., Perfusion MRI in treatment evaluation of glioblastomas: Clinical relevance of current and future techniques. Journal of Magnetic Resonance Imaging, 2019. 49(1): p. 11–22.

3. Pantel, A.R., et al., PET Imaging of Metabolism, Perfusion, and Hypoxia: FDG and Beyond. The Cancer Journal, 2024. 30(3).

4. Li, E.J., et al., Metabolic Imaging and Molecular Signatures Reveal Metabolic Phenotypes in Recurrent Glioblastoma. Journal of Nuclear Medicine, 2026. 67(6): p. 849–856.

5. Lortie, M., et al., Quantification of myocardial blood flow with 82Rb dynamic PET imaging. European Journal of Nuclear Medicine and Molecular Imaging, 2007. 34(11): p. 1765–1774.

6. Sciagrà, R., et al., EANM procedural guidelines for PET/CT quantitative myocardial perfusion imaging. European Journal of Nuclear Medicine and Molecular Imaging, 2021. 48(4): p. 1040–1069.

7. Hygino da Cruz, L.C., et al., Pseudoprogression and Pseudoresponse: Imaging Challenges in the Assessment of Posttreatment Glioma. American Journal of Neuroradiology, 2011. 32(11): p. 1978–1985.

8. Sahani, D.V., et al., Assessing tumor perfusion and treatment response in rectal cancer with multisection CT: Initial observations. Radiology, 2005. 234(3): p. 785–792.

9. Carson, R.E., M. Naganawa, and J.-D. Gallezot, PET Quantification and Kinetic Analysis, in Molecular Imaging of Neurodegenerative Disorders. 2023, Springer International Publishing. p. 183–194.

10. Watabe, H., Compartmental Modeling in PET Kinetics, in Basic Science of PET Imaging. 2017, Springer International Publishing. p. 323–352.

11. Koonce, J.D., et al., Accuracy of dual-energy computed tomography for the measurement of iodine concentration using cardiac CT protocols: validation in a phantom model. Eur Radiol, 2014. 24(2): p. 512–8.

12. Wang, Y., et al., Total-Body PET Kinetic Modeling and Potential Opportunities Using Deep Learning. PET Clin, 2021. 16(4): p. 613–625.

13. Shapira, N., et al., Quantitative positron emission tomography imaging in the presence of iodinated contrast media using electron density quantifications from dual-energy computed tomography. Medical Physics, 2021. 48(1): p. 273–286.

14. Scherer, K., et al., Dynamic Quantitative Iodine Myocardial Perfusion Imaging with Dual-Layer CT using a Porcine Model. Scientific Reports, 2019. 9(1): p. 16046.

15. Hoffman, E.J., et al., 3-D phantom to simulate cerebral blood flow and metabolic images for PET. IEEE Transactions on Nuclear Science, 1990. 37(2): p. 616–620.

16. Zhao, B., Lung Phantom. 2015, The Cancer Imaging Archive.

17. Gulliksrud, K., C. Stokke, and A.C. Trægde Martinsen, How to measure CT image quality: Variations in CT-numbers, uniformity and low contrast resolution for a CT quality assurance phantom. Physica Medica, 2014. 30(4): p. 521–526.

18. Gabrani-Juma, H., et al., Validation of a Multimodality Flow Phantom and Its Application for Assessment of Dynamic SPECT and PET Technologies. IEEE Transactions on Medical Imaging, 2017. 36(1): p. 132–141.

19. Krakovich, A., et al., A new cardiac phantom for dynamic SPECT. Journal of Nuclear Cardiology, 2021. 28(5): p. 2299–2309.

20. Black, D.G., et al., Design of an anthropomorphic PET phantom with elastic lungs and respiration modeling. Medical Physics, 2021. 48(8): p. 4205–4217.

21. Kamphuis, M.E., et al., A Multimodality Myocardial Perfusion Phantom: Initial Quantitative Imaging Results. Bioengineering, 2022. 9(9): p. 436.

22. Driscoll, B., H. Keller, and C. Coolens, Development of a dynamic flow imaging phantom for dynamic contrast-enhanced CT. Med Phys, 2011. 38(8): p. 4866–80.

23. Hannuksela, M., S. Lundqvist, and B. Carlberg, Thoracic aorta – dilated or not? Scandinavian Cardiovascular Journal, 2006. 40(3): p. 175–178.

24. Dai, B., et al., Performance evaluation of the PennPET explorer with expanded axial coverage. Physics in Medicine and Biology, 2023. 68(9).

25. Peristaltic Pump Tubing Specification. Aurora Pro Scientific LLC: Midland Park, NJ.

26. Li, E.J., et al., Efficient Delay Correction for Total-Body PET Kinetic Modeling Using Pulse Timing Methods. J Nucl Med, 2022. 63(8): p. 1266–1273.

27. Li, E.J., et al., Total-Body Perfusion Imaging with [11C]-Butanol. J Nucl Med, 2023.

